# [^18^F]ROStrace detects oxidative stress *in vivo* and predicts progression of Alzheimer’s disease pathology in APP/PS1 mice

**DOI:** 10.1101/2022.04.05.487092

**Authors:** Chia-Ju Hsieh, Catherine Hou, Yi Zhu, Ji Youn Lee, Neha Kohli, Evan Gallagher, Kuiying Xu, Hsiaoju Lee, Shihong Li, Meagan J. McManus, Robert H. Mach

**Affiliations:** Department of Radiology, Perelman School of Medicine, University of Pennsylvania, Philadelphia, Pennsylvania 19104, USA; Department of Anesthesiology and Critical Care Medicine, The Children’s Hospital of Philadelphia, Philadelphia, PA, USA; Center for Mitochondrial and Epigenomic Medicine, The Children’s Hospital of Philadelphia, Philadelphia, Pennsylvania, USA

**Keywords:** Oxidative Stress, Positron Emission Tomography, ROStrace, Alzheimer’s Disease, Neuroinflammation, Neurodegeneration

## Abstract

**Purpose:** Oxidative stress is implicated in the pathogenesis of the most common neurodegenerative diseases, such as Alzheimer’s disease (AD). However, tracking oxidative stress in the brain has proven difficult and impeded its use as a biomarker. Herein, we investigate the utility of a novel positron emission tomography (PET) tracer, [^18^F]ROStrace, as a biomarker of oxidative stress throughout the course of AD in the well-established APP/PS1 double mutant mouse model.

**Methods:** PET imaging studies were conducted in wild-type (WT) and APP/PS1 mice at 3 different time points, representing early (5 mo.), middle (10 mo.), and advanced (16 mo.) life (n = 6-12, per sex). Semi-quantitation SUVRs of the plateau phase (40-60min post-injection; SUVR_40-60_) of ten brain subregions were designated by the Mirrione atlas and analyzed by Pmod. Statistical parametric mapping (SPM) was used to distinguish brain regions with elevated ROS in APP/PS1 relative to WT in both sexes. The PET studies were validated by *ex vivo* autoradiography and immunofluorescence with the parent compound, dihydroethidium.

**Results:** [^18^F]ROStrace retention was increased in the APP/PS1 brain compared to age-matched controls by 10 mo. of age (p < 0.0001), and preceded the accumulation of oxidative damage in APP/PS1 neurons at 16mo. (p < 0.005). [^18^F]ROStrace retention and oxidative damages were higher and occurred earlier in female APP/PS1 mice as measured by PET (p < 0.001), autoradiography and immunohistochemistry (p < 0.05). [^18^F]ROStrace differences emerged mid-life, temporally and spatially correlating with increased Aβ burden (r^2^ = 0.36; p = 0.0003), which was also greatest in the female brain (p < 0.001).

**Conclusions:** [^18^F]ROStrace identifies increased oxidative stress and neuroinflammation in APP/PS1 female mice, concurrent with increased amyloid burden mid-life. Differences in oxidative stress during this crucial time may partially explain the sexual dimorphism in AD. [^18^F]ROStrace may provide a long-awaited tool to stratify at-risk patients who may benefit from antioxidant therapy prior to irreparable neurodegeneration.

## Introduction

Alzheimer’s disease (AD) is the most common neurodegenerative disease worldwide. Over the past 60 years, life expectancy in the United States has increased by almost a decade, tripling the number of people over the age of 65 to a record high of 50 million. One in ten of these individuals is diagnosed with AD, and two-thirds of those are women [1]. In the absence of preventative therapeutics or a cure, the number of AD sufferers will reach approximately 130 million worldwide by 2050 [2].

AD pathogenesis is estimated to begin decades before symptoms emerge [3, 4]. Biomarkers that track the early pathogenic changes may provide an optimal therapeutic window by identifying those at risk of AD before irreversible brain damage and cognitive impairment occurs. Evidence from preclinical models and at-risk patients suggests the earliest biochemical changes in the AD brain involve metabolic decline [5, 6], oxidative stress [7-12], inflammation [13], and aberrant processing of amyloid precursor protein (APP) and tau [14-17]. Interestingly, each of these key changes has been linked to mitochondrial dysfunction and the production of reactive oxygen species (ROS) [18, 19]. Mitochondrial function begins to decline during middle age [20], which may underlie the maternally-inherited pattern of glucose hypometabolism in the at-risk AD brain [6, 19, 21, 22]. Impaired mitochondria use ROS to alert the cellular environment of impending bioenergetic stress [23]. ROS refers to a family of partially reduced oxygen species, such as O_2_^•-^, HO^•^, H_2_O_2_, NO, and ONOO^-^, that possess highly reactive properties. ROS directly activate microglia, the resident macrophages of the brain. Activated microglia in turn produce more ROS via NADPH oxidase (NOX) and nitric oxide synthase (NOS) [24, 25]. When the resulting rise in ROS production overwhelms the modest level of endogenous antioxidants in the aging brain, oxidative stress ensues. Oxidative stress modifies the enzymes responsible for pathogenic processing of amyloid precursor protein (APP) and tau, leading to Aβ and tau accumulation [18]. Aβ and tau, along with oxidized lipids and DNA, can shift microglia into a chronic, ROS-generating phenotype [23, 25-32], thereby propelling the at-risk AD brain into a pro-inflammatory state that is neurotoxic. The primary ROS produced by mitochondria and activated microglia is superoxide (O_2_^•-^). Thus, O_2_^•-^ provides a signal of mitochondrial and immune (mito-immune) stress, which may be used to track neuroinflammation throughout the course of AD.

Our group has pioneered the development of positron emission tomography (PET) radiotracers for imaging oxidative stress *in vivo* [33, 34]. The parent compound for radiotracer development is dihydroethidium (DHE), which is primarily oxidized by O_2_^•-^ to ethidium and trapped in tissues [35]. Prior validation of [^18^F]ROStrace demonstrates that it behaves similarly to DHE.

Importantly, only the neutral species of [^18^F]ROStrace crosses the blood brain barrier. Once in the brain, if [^18^F]ROStrace is oxidized, [^18^F]*ox*-ROStrace becomes trapped. This trapping mechanism results in increased [^18^F]ROStrace retention in conditions of increased oxidative stress and neuroinflammation [34].

The two major goals of the current study were to determine if [^18^F]ROStrace is sensitive enough 1) to detect increased oxidative stress in a preclinical model of AD and 2) to predict the accelerated AD pathology in the female brain. Increased Aβ is reported to be the earliest neuropathological change in the at-risk AD brain, and the presence of Aβ is currently required for AD diagnosis [36, 37]. We therefore analyzed the relationship of [^18^F]ROStrace and Aβ in the well-characterized, double transgenic APP_SWE_/PS1dE9 (APP/PS1) mouse model. Much like AD patients, Aβ plaques, neuroinflammation, and cognitive defects are more prevalent in female compared to male APP/PS1 mice [38-40]. Due to the established role of oxidative stress in Aβ pathology, we hypothesized that [^18^F]ROStrace retention would correlate with increased Aβ burden in female APP/PS1 mice. To test this hypothesis, a series of PET imaging studies were conducted in wild-type (WT) and APP/PS1 mice of both sexes at 3 different time points, representing early (5 mo.), middle (10 mo.), and advanced (16 mo.) life. The results of our study show that [^18^F]ROStrace is increased in the APP/PS1 brain compared to WT. Further, [^18^F]ROStrace retention is higher and occurs earlier in female APP/PS1 mice. Thus, female AD mice are more prone to oxidative stress than males, and this difference emerges with increased Aβ burden mid-life. Our results suggest that differences in oxidative stress during this crucial time may partially explain the sexual dimorphism in AD.

## Results

### [^18^F]ROStrace retention is increased in APP/PS1 mice in vivo and reveals a female sex bias

Oxidative stress is a driver of AD pathology in APP/PS1 mice [41, 42]. To determine the sensitivity of [^18^F]ROStrace to detect increased oxidative stress during the course of AD, a series of micro-PET imaging studies were performed in WT and APP/PS1 mice at 5 mo., 10 mo., and 16 mo. of age (online Fig. SI 1). Due to the lack of blood sample collection to perform absolute quantitation on [^18^F]ROStrace PET images and differences in body weight between female and male (Fig. 1d), we utilized a semi-quantitative approach based on normalization to a reference region. However, since ROS produced by mitochondria or activated microglia could be throughout the brain, there is no true reference region for quantitating [^18^F]ROStrace images. Studies of TSPO neuroinflammatory PET imaging agents have suggested using a pseudo-reference region, which is a region of interest (ROI) that has no difference in SUVs among the study groups, to increase the sensitivity of the PET image analysis [43, 44]. In the current study, the periaqueductal gray (PAG) was the ROI showing no statistical differences in the [^18^F]ROStrace SUVs among WT and APP/PS1 in both genders at all ages (Fig. 1e). Therefore, this ROI was selected as the pseudo-reference region for [^18^F]ROStrace SUVR measurement in this preclinical AD study.

**Fig. 1.**
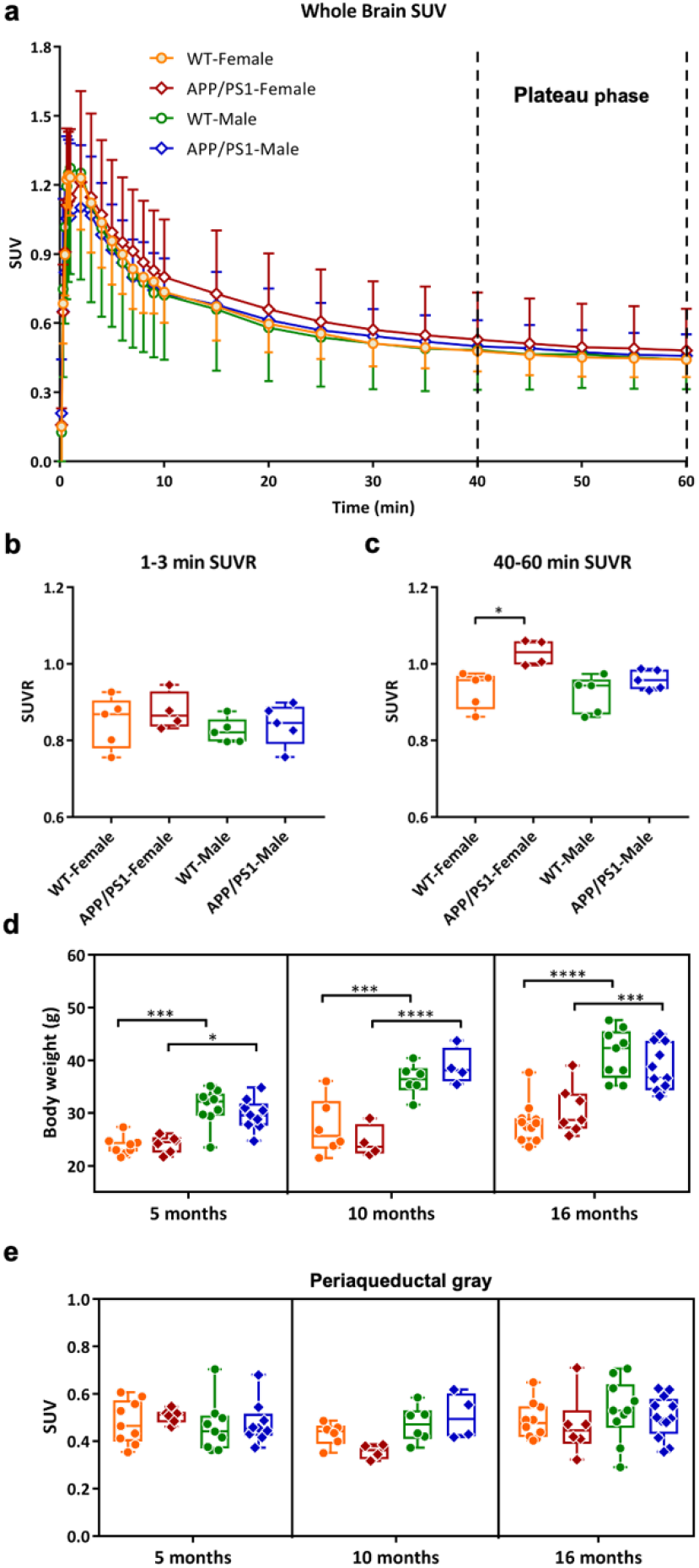
Validation of the plateau phase and pseudo-reference region for ox[^18^F]ROStace PET analysis. Micro-PET data analysis validates that oxidative stress, not blood flow, contribute the [^18^F]ROStace signals in the animal brain. Time activity curves in SUV of whole brain (**a**) in WT female (n=5), WT male (n=5), APP/PS1 female (n=4), and APP/PS1 male (n=5) mice at 16 mo. of age. Data is shown as mean ± standard deviation. Comparison of the average [^18^F]ROStrace SUVR from 9 VOIs at 16 mo. old animals in (**b**) perfusion phase (1-3 min) and (**c**) plateau phase (40-60 min). (WT groups: n = 5 per gender, and APP/PS1 group: n = 4 for female, and n = 5 for male) The statistical significance of p value calculated by one-way ANOVA (* p < 0.05). Comparison of (**d**) Body weight (g) and (**e**) Periaqueductal gray (PAG) SUV from 40-60 min. 5 mo. old: APP/PS1 group: n = 6 for female, and n = 9 for male; WT group: n = 9 per gender,10 mo. old: APP/PS1 group: n = 4 per gender; WT group: n = 6 per gender, and16 mo. old: APP/PS1 group: n = 6 for female, and n = 12 for male; WT group: n = 10 per gender. The statistical significance of p value calculated by two-way ANOVA: **** p < 0.0001, *** p < 0.001, * p < 0.05

First, whole brain time-activity curves were generated from dynamic scans taken 0-60 min post-injection of [^18^F]ROStrace (Fig. 1a). As previously reported [34], [^18^F]ROStrace reached peak uptake in brain within 2 min, followed by a washout period for approximately 20 min, and then remained steady in a plateau phase throughout the remainder of the scan. There were no differences in SUVRs during the perfusion phase (1-3 min; Fig. 1b), indicating that [^18^F]ROStrace levels during the plateau phase, which represent oxidized [^18^F]ROStrace [34], cannot be attributed to differences in blood flow (Fig. 1c).

Next, regional semi-quantitation SUVRs of the plateau phase (40-60min post-injection; SUVR_40-60_) were calculated for WT and APP/PS1 mice at the three different time points (Fig 2). Representative microPET images of region-specific [^18^F]ROStrace retention for each strain, age, and sex are shown in Fig. 2a. To confirm our microPET results, [^18^F]ROStrace activity was also measured using ex vivo autoradiography (ARG) (Fig. 2a). [^18^F]ROStrace retention was indistinguishable amongst all brain regions of the APP/PS1 and WT mice at the earliest time point of 5 mo. (Fig. 2b). [^18^F]ROStrace retention was higher in the cortex and cerebellum of the APP/PS1 females compared to APP/PS1 males (p < 0.05; Table S1). This sex-based difference was amplified at 10 mo. of age, where APP/PS1 females demonstrated increased [^18^F]ROStrace retention over that of APP/PS1 males in every brain region except the midbrain and brainstem (Fig. 2c). Compared to WT mice, [^18^F]ROStrace retention was increased in the cortex (p = 0.0002), striatum (p = 0.0116), hippocampus (p = 0.0336), and amygdala (p = 0.0010) of female APP/PPS1, whereas APP/PS1 males remained indistinguishable from WT (Fig. 2c).

**Fig. 2.**
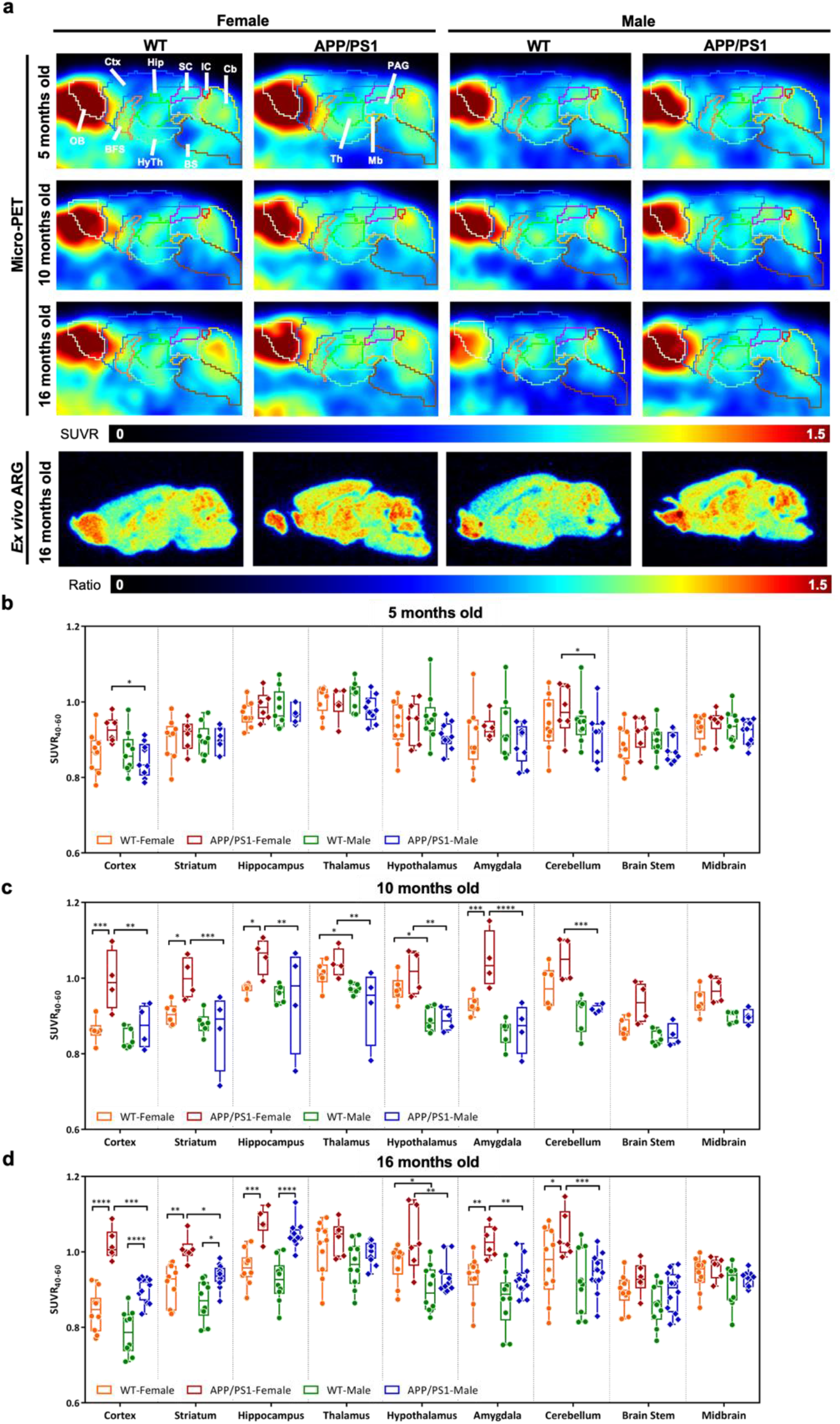
*In vivo* [^18^F]ROStrace retention is increased in the APP/PS1 model of AD and demonstrates a female sex bias. (**a**) Sagittal view of [^18^F]ROStrace SUVR_40-60_ PET images collected from both sexes of WT and APP/PS1 mice at 5 mo., 10 mo., and 16 mo. Brain subregions derived from Mirrione mouse brain atlas. Ctx: cortex, OB; olfactory bulb, BFS: basal forebrain septum, Hip: hippocampus, HyTh: hypothalamus, SC: superior colliculi, IC: inferior colliculi, Cb: cerebellum, BS: brain stem, Th: thalamus, PAG: Periaqueductal gray, and Mb: midbrain. Region-specific differences detected by [^18^F]ROStrace PET were confirmed by *ex vivo* autoradiography (ARG) from the same animals at 16 mo. (**b-d**) Regional differences in [^18^F]ROStrace uptake from 9 VOIs presented as standard uptake value ratios from 40-60min post-injection (SUVR_40-60_) in WT and APP/PS1 mice at (**b**) 5 mo. old: APP/PS1 group: n = 6 for female, and n = 9 for male; WT group: n = 9 per gender, (**c**) 10 mo. old: APP/PS1 group: n = 4 per gender; WT group: n = 6 per gender, and (**d**) 16 mo. old: APP/PS1 group: n = 6 for female, and n = 12 for male; WT group: n = 10 per gender. The statistical significance of p value calculated by two-way ANOVA (*p < 0.05, **p < 0.01, ***p < 0.001, ****p < 0.0001).

The pattern of increased [^18^F]ROStrace retention in female APP/PS1 *vs*. WT was similar at 10 and 16 mo. of age, but more significant in the cortex (p < 0.0001), striatum (p = 0.0034), hippocampus (p = 0.0001), amygdala (p = 0.0031), and cerebellum (p = 0.0257) (Online Table S1). Finally, at 16mo. of age, [^18^F]ROStrace retention increased in APP/PS1 male mice. Higher SUVR_40-60_ levels were observed in APP/PS1 *vs*. WT male in cortex (p < 0.0001), striatum (p = 0.0217), and hippocampus (p < 0.0001) (Fig. 2d). Notably, the only region with differential [^18^F]ROStrace retention in female *vs*. male WT mice was the hypothalamus from mid- (p = 0.0156) to advanced (p = 0.0165) life.

In order to understand the effect of aging on retention of [^18^F]ROStrace in brain tissue, SUVR_40-60_ measurements were compared within each strain over time using a two-way ANOVA. The statistical comparisons show a decrease in several regions in male WT mice from 5 to 16 mo., but [^18^F]ROStrace remained stable in most regions of the WT female brain over time (Online Table S1). Conversely, [^18^F]ROStrace retention increased over time in mice with APP/PS1 mutations, regardless of sex. From 5 to 16 mo., [^18^F]ROStrace retention increased in the hippocampus (p = 0.0055), striatum (p = 0.0023), cortex (p = 0.0050), hypothalamus (p = 0.0204), and amygdala (p = 0.0157) of APP/PS1 females, but only the hippocampus (p = 0.0018) and cortex (p = 0.0296) of APP/PS1 males.

### [^18^F]ROStrace retention correlates with APP/PS1 amyloid burden

ROS induce Aβ-generating β- and γ-secretases [45], while inhibiting Aβ-clearing enzymes [46]. The resulting Aβ aggregates are associated with activated microglia and oxidative damage in the AD brain. Therefore, we hypothesized that [^18^F]ROStrace levels would correlate with Aβ burden in APP/PS1 mice in a region-specific manner. To test this hypothesis, statistical parametric mapping (SPM) was first used to distinguish brain regions with elevated ROS in APP/PS1 relative to WT in both sexes (Fig. 3a). The SPM results show elevation of [^18^F]ROStrace retention in APP/PS1 female mice at 10 mo. of age in hippocampus and cortex. By 16 mo., [^18^F]ROStrace levels were augmented and extended to broader areas of both regions (Fig. 3a). In APP/PS1 males, higher [^18^F]ROStrace was also observed in the hippocampus and cortex, but only at the most advanced age.

**Fig. 3.**
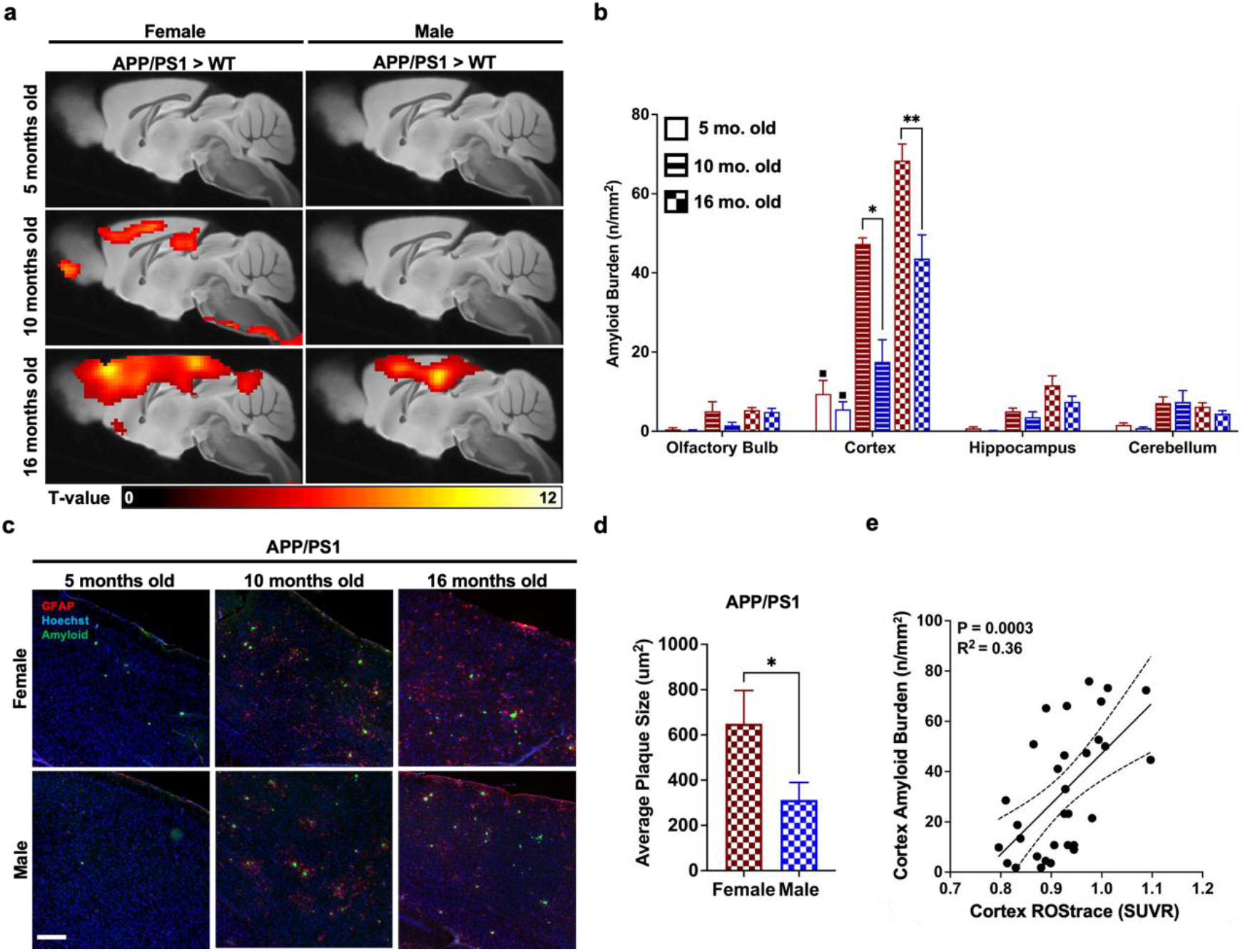
Elevated [^18^F]ROStrace retention in the APP/PS1 brain maps with spatial distribution of amyloid deposits and positively correlates with increased amyloid burden in the cortex of aging APP/PS1 mouse brain. (**a**) Statistical parametric mapping (SPM) analysis showing significant differences (uncorrected p < 0.005 and an extent cluster threshold = 50) in [^18^F]ROStrace retention per sex in 5 mo. (top row), 10 mo. (middle row), and 16 mo. (bottom row) APP/PS1 mice *vs*. WT. (**b**) Quantification of amyloid burden in 4 different brain regions of the APP/PS1 female (red) and male (blue) mouse brain. 5 mo. old APP/PS1 group: n = 9 for male, n = 5 for female, 10 mo. old APP/PS1 group: n = 4/gender. 16 mo. old APP/PS1 group: n = 8 for male, n = 5 for female. The statistical significance calculated by two-way ANOVA. * p < 0.05, ** p < 0.005. ^▪^ p < 0.0001, relative to olfactory bulb, hippocampus and cerebellum at 5mo. old APP/PS1 mouse. Values represent mean ± SEM. (**c**) Images of the cortex region of APP/PS1 mice 5 – 16 mo. of age stained with anti-GFAP (red), Anti-beta Amyloid 1-42 antibodies (green) and Hoechst (blue). Scale bar = 150 μm. (**d**). Average size of amyloid plaques in the cortex region of brains collected from 16 mo. old APP/PS1 mice, n = 8 for male, n = 5 for female. The statistical significance of p value calculated by unpaired *t*-test. Values represent mean ± SEM. * p < 0.05 (**e**) Linear regression analysis of cortical amyloid burden (n/mm^2^) and [^18^F]ROStrace SUVR_40-60_.

Next, the brains from 5-, 10- and 16-mo. old APP/PS1 mice were processed with anti-amyloid and GFAP antibodies. During pro-inflammatory conditions like trauma and amyloidosis, astrocytes assist in neuronal repair by creating a barrier to confine toxic elements and pave the way for microglial phagocytosis [47]. Our co-staining results confirm that amyloid plaques were closely surrounded by GFAP-positive astrocytes in the APP/PS1 brain. The representative images in online Fig. SI 2 show that very few Aβ aggregates were visible at 5 mo. Of age in either sex, and these were predominately found in the olfactory bulb and the cortex. Female APP/PS1 mice also had sparse, small Aβ aggregates in the cerebellum at 5 mo. By 10 mo. of age, Aβ aggregates were detected throughout all four brain regions (olfactory bulb, cortex, hippocampus and cerebellum), but were most prevalent in the cortex and hippocampus. A similar trend of Aβ distribution was seen at 16 mo.

To further investigate the relationship between regional amyloid burden and [^18^F]ROStrace, low magnification images of sagittal brain sections were acquired from APP/PS1 mice in each age group and the number of amyloid aggregates quantified in 4 key regions (Fig. 2b and online Fig SI 2). The results show that the cortex is not only the first brain region where significant amyloid plaques appeared (p < 0.0001 for cortex *vs*. olfactory bulb, hippocampus, and cerebellum at 5 mo.), but also the region that bore the heaviest amyloid burden with increasing age (Fig. 3b, c). Amyloid burden in the cortex was higher in APP/PS1 females than in age-matched males, which became significant at 10 mo. (Fig. 3b). The size of amyloid plaques was also largest in the cortex of APP/PS1 female mice at 16 mo. (Fig. 3d). These results confirm previous reports of increased Aβ burden in APP/PS1 females [38, 39]. Interestingly, the cortex was also the first region to show increased [^18^F]ROStrace levels in both sexes of APP/PS1 mice, and [^18^F]ROStrace remained consistently higher in the APP/PS1 female *vs*. male cortex over time. To evaluate the significance of this association, linear regression analysis was performed for amyloid burden *vs*. [^18^F]ROStrace SUVR_40-60_ in the cortex of APP/PS1 mice, which revealed a positive correlation between amyloid burden and [^18^F]ROStrace (p = 0.0003, R^2^ = 0.36; Fig. 3e). Collectively, these results highlight the sensitivity of [^18^F]ROStrace to predict AD progression.

### Validation of increased oxidative stress in the APP/PS1 brain

We further characterized the oxidative stress detected by [^18^F]ROStrace *in vivo* by analyzing the same brains with established fluorescent detection methods. Following the PET scan, mice were injected with the parent compound, DHE (5mg/kg, i.p.) and euthanized 30min later. To estimate the cellular source of the [^18^F]ROStrace signal, colocalization between DHE and three major brain cell types was then determined using cell-type specific markers for neurons (NeuN), microglia (IBA-1) and astrocytes (GFAP) (Fig. 4). The fluorescent images revealed that DHE was localized to the cytoplasm of neurons in the APP/PS1 brain only (Fig. 4a, b), which may represent Aβ-induced mitochondrial ROS [12, 48]. DHE was also frequently found in aggregated microglia in the APP/PS1 brain. These aggregated microglia surrounded amyloid plaques and had an amoeboid shape, with a large cell body and relatively few, short processes, resembling an activated morphology (Fig. 4d). In contrast, DHE was rarely detected in the microglia of age-matched WT brain (Fig. 4c). Lastly, DHE signal showed no detectable colocalization with GFAP, indicating that astrocytes have minimal involvement in the ROS signal detected by DHE (Fig. 4e, f), and presumably [^18^F]ROStrace, in both WT and APP/PS1 animals.

**Fig. 4.**
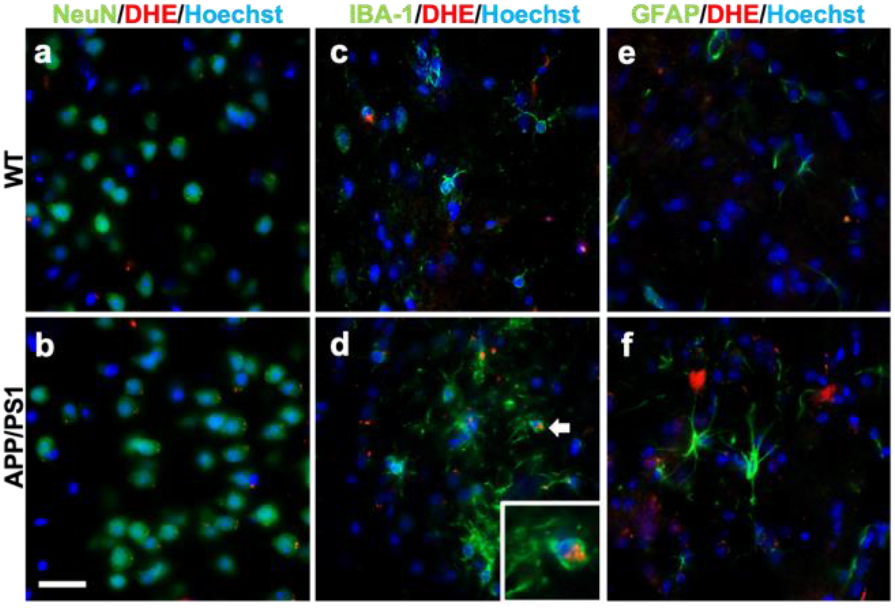
DHE colocalizes with microglia and neurons, but not astrocytes, in the APP/PS1 brain. Representative confocal micrographs showing colocalization of (**a-b**) oxidized DHE (red) with neurons (NeuN; green) and (**c-d**) microglia (Iba1; green) in WT and APP/PS1 cortex. (**e-f**) Oxidized DHE did not colocalize with astrocytes (GFAP, green). The Inlet is a magnified region of that indicated by the white arrow. Scale bar = 20 μm.

One of the key mediators of ROS-induced neurotoxicity in AD is peroxynitrite, (ONOO^-^) [49, 50]. ONOO^-^ is a powerful oxidant produced by O_2_·- and nitric oxide (·NO), that permanently alters proteins via nitration of tyrosine residues. To determine the relationship between O_2_^•-^ detected by [^18^F]ROStrace and downstream oxidative damage, we quantified levels of 3-nitrotryosine (3NT) in age-matched WT and APP/PS1 mouse brains using IHC. A gradual elevation of 3NT was observed in the cortex of aging APP/PS1 mouse brain (Online Fig. SI 3), which only became significant at 16 mo. of age (Fig. 5a). As expected, APP/PS1 females suffered the most extensive oxidative damage, evidenced by the highest number of 3NT-positive cells (Fig. 5b).

**Fig. 5.**
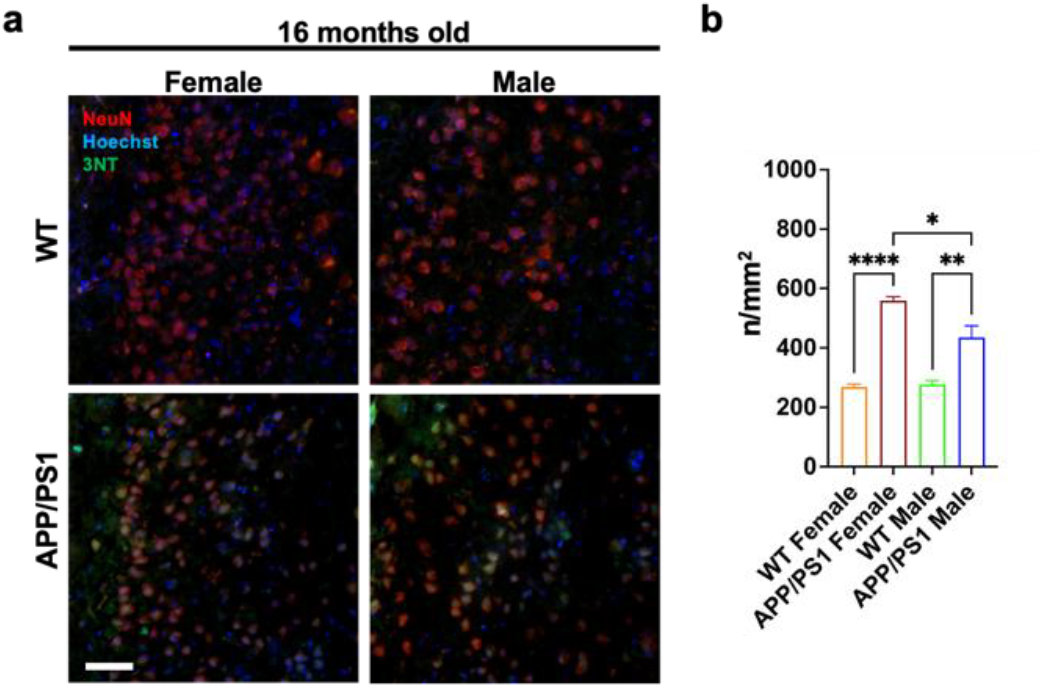
Oxidative damage is highest in the female APP/PS1 at advanced age. (**a**) Representative confocal images showing increased oxidative damage in neurons (NeuN; red) of APP/PS1 mice detected by 3-Nitrotyrosine (3NT; green) adducts in the cortex at 16 mo. of age. Scale bar = 50 μm. (**b**) Quantification of 3NT positive cells in the cortex of age-match WT and APP/PS1 mouse brain tissues, n = 4 per group. The statistical significance of p value calculated by one-way ANOVA. Values represent mean ± SEM, * p < 0.05, ** p < 0.005, **** p < 0.0001.

## Discussion

The preclinical studies described herein suggest that oxidative stress may contribute to the increased risk of AD in women. Using the novel superoxide sensitive PET tracer, [^18^F]ROStrace, we detected increased oxidative stress in the female AD brain *in vivo*, which coincided with increased Aβ and preceded the accumulation of neuronal damage. Increased Aβ is one of the earliest neuropathological changes in the at-risk AD brain, and is required for AD diagnosis [36, 37]. However, the disconnect between Aβ and cognitive performance presents a major limitation and suggests additional biomarkers are needed to predict the progression of neurodegeneration and gauge therapeutic efficacy throughout the course of disease [51]. Our results suggest [^18^F]ROStrace is a promising tool to bridge this gap.

Mitochondrial dysfunction and microglial activation occur early in the neurodegeneration process, leading to a collective increase in ROS production and oxidative stress that continues through the course of AD. Markers of oxidative damage have been reported in mild cognitive impairment and early AD, but longitudinal analysis has thus far been limited to biofluids. Herein, we confirm the early and persistent role of oxidative stress in the brain using [^18^F]ROStrace PET. We found that the fluorescent parent compound of [^18^F]ROStrace, DHE, colocalized to neurons and microglia in the AD brain. These observations are in line with prior studies describing increased production of O_2_^•-^ by neuronal mitochondria and activated microglia in response to Aβ [12, 32, 52, 53]. In addition to O_2_^•-^, Aβ also induces nitric oxide (^•^NO) in both mitochondria and microglia [54, 55]. ^•^NO is the only molecule that can out-compete superoxide dismutase for O_2_^•-^, reacting with O_2_^•-^ at an estimated rate of 4-16 ×10^9^ M ^-1^s ^-1^ to form peroxynitrite, ONOO^-^ [56, 57]. ONOO^-^ permanently modifies proteins by nitrating tyrosine residues, which can thus be used as a molecular footprint of oxidative damage [58]. Using 3-nitrotyrosine, we found oxidative damage increased over time in the AD brain, as expected [49, 50], but was highest in females. Notably, the increase in [^18^F]ROStrace retention preceded oxidative damage in neurons, suggesting [^18^F]ROStrace may serve as a predictive biomarker of downstream neuronal demise.

The mito-immune shifts that occur in the female brain due to estrogen changes mid-life are thought to be a major contributor to the increased risk of females to AD [59]. We selected 3 time-points for [^18^F]ROStrace imaging, that for female mice, represent key states which closely resemble the human condition during premenopause (5 mo.), perimenopause (10 mo.), and postmenopause (16 mo.) [60-62]. Interestingly, the only region with significantly different [^18^F]ROStrace retention in WT females and males was the hypothalamus. The hypothalamus plays an important role of neuroendocrine function, especially in regulating estrogen in females [63-66]. The hypothalamic increase in [^18^F]ROStrace retention in females began at 10 mo., which represents the mid-point of the proposed perimenopausal phase (7-12 mo.) in mice [61, 62]. Perimenopause describes the mid-life transition to reproductive senescence. Perimenopause also represents a critical transition period in the aging female brain, requiring metabolic adaptation to the decline in estrogen-dependent bioenergetics [63]. Most women maintain the neurological resilience to transition through perimenopause without long-term consequences. However, for those at risk of AD, the neurological challenges involved in perimenopause may overwhelm the system and tip the scales towards neurodegeneration. Our results suggest that oxidative stress is a key indicator of this proposed tipping point.

Estrogen regulates mitochondrial biogenesis [67, 68], endogenous antioxidants [69], and microglial activation [40, 70]. Thus, in an environment of increased Aβ, such as the APP/PS1 brain, a decline in estrogen signaling would act as a second hit on mitochondrial function, leading to increased ROS production [71]. Increased mitochondrial ROS can drive the conversion of microglia into a pro-inflammatory phenotype by impairing oxidative phosphorylation, inducing mitochondrial fission, activating nuclear factor kappa B (NF-kB), and stimulating the metabolic switch to glycolysis [72]. Pro-inflammatory microglia amplify ROS production, which can become neurotoxic. Microglia in the female AD brain are predominately pro-inflammatory and less phagocytic than that of males, leading to increased amyloid burden and oxidative stress [40]. In preclinical models, oxidative stress induces every established pathological hallmark of AD [18]. Therefore, increased oxidative stress in the brain during mid-life, when AD-related neurodegeneration is proposed to begin, may serve as a predictive biomarker of AD pathogenesis. Future studies are required to determine the underlying mechanisms responsible for increased oxidative stress in females, and whether beta-estradiol or other antioxidant therapy administered during this critical, early phase may prevent oxidative stress and slow AD progression in at risk patients.

## Materials and Methods

### Animal models and experimental scheme

All animal experiments in this study were performed under protocols approved by the University of Pennsylvania Institutional Animal Care and Use Committee (IACUC). The double transgenic APP_SWE_/PS1dE9 (APP/PS1) mice expressing Mo/HuAPP695swe (chimeric mouse and human amyloid precursor protein) and PS1-dE9 (mutant human presenilin 1), and the littermate control C57Bl/6J (WT; wild type) were purchased from Mutant Mouse Resource and Research Center (MMRRC). A total of 91 mice including 41 APP/PS1 (5 mo.: 6 female and 9 male; 10 mo.: 4 female and 4 male; 16 mo.: 6 female and 12 male) and 50 WT (5 mo.: 9 female and 9 male; 10 mo.: 6 female and 6 male; 16 mo.: 10 female and 10 male) were used in this study. The general experimental scheme of this study is shown in Fig S1. Briefly, animals selected from each age group received 200∼300 μCi of [^18^F]ROStrace via tail vein injection. After a 20 mins (40-60 min post-injection) or a one hour (0-60 min post-injection) dynamic PET scan, a dose of 5 mg/kg DHE (Sigma-Aldrich, St. Louis, MO, USA) was injected to the animals. The animals were sacrificed 30 mins after DHE injection and brains were extracted for subsequent experiments.

### Preparation of [^18^F]ROStrace

The radiosynthesis of [^18^F]ROStrace was prepared as previously described [34]. Briefly, [^18^F]ROStrace was accomplished on all-in-One module (Trasis, Belgium) with full automation. The final product was diluted with 0.6 mL of ethanol, 0.1% ascorbic acid and 6 mL normal saline, and filtered by 0.2 µM nylon filter. The radiochemical yield was 4∼20%, and the specific activity was 2000 Ci/mmol.

### Micro-PET imaging

PET imaging was performed on the β-Cube PET scanner (Molecubes, Ghent, Belgium). The body weight of each animal was measured before the PET scan (Fig S2A). Animals were anesthetized with 1-2% isoflurane, a tail vein catheter was placed for PET tracer administration, and the animal was placed on the scanner bed. Dynamic scans of 0-60 min or 40-60 min post-injection were acquired after injection of 200-300 μCi of [^18^F]ROStrace. After completion of the PET scan, the animal was transferred to the X-Cube CT scanner (Molecubes, Ghent, Belgium) and a general-purpose CT scan was acquired for anatomical reference and attenuation correction. PET images were reconstructed with a matrix size of 192 × 192 × 384, and a voxel size of 0.4 × 0.4 × 0.4 mm with frame lengths of 6 × 10 sec, 9 × 60 sec, and 10 × 300 sec for 60 minutes dynamic scans or 4 × 300 sec for 20 minutes scans. All corrections were applied using a manufacturer supplied reconstruction program. CT images were reconstructed with a matrix size of 200 × 200 × 550, and a voxel size of 0.2 × 0.2 × 0.2 mm with a manufacturer supplied reconstruction program.

### Micro-PET image analysis

All the [^18^F]ROStrace micro-PET/CT imaging data were processed and analyzed by using Pmod software (version 3.7, PMOD Technologies Ltd., Zurich, Switzerland). Rigid body matching was manually performed on individual micro-CT images to co-register to the Mirrione mouse MR-T2 weighted brain template [1]. Then, the resulting transformation parameters were applied to the corresponding micro-PET image. Ten volumes of interest (VOIs) including cortex, thalamus, cerebellum, hypothalamus, brain stem, periaqueductal gray, striatum, hippocampus, amygdala, and midbrain were selected from the Mirrione atlas [73]. Due to the lack of the blood samples to perform absolute quantitation and a true reference region to semi-quantitate neuroinflammation images, a pseudo-reference region [44] is needed for [^18^F]ROStrace neuroinflammation imaging evaluation. Therefore, the periaqueductal gray, with no significant difference in standardized retention values (SUVs; Fig S2B) among the different animal groups, was selected as the pseudo-reference region for calculating the SUV ratio (SUVR) for each VOI. The 1-3 min SUVR and the 40-60 min SUVR (SUVR_40-60_) were extracted from all the VOIs for the perfusion and the late phase comparison from the nineteen 16 mo. old animals (5 WT female, 5 WT male, 4 APP/PS1 female, and 5 APP/PS1 male) with 1 hour full dynamic [^18^F]ROStrace imaging.

Voxel-wise analyses were performed by using Statistical Parametric Mapping 12 (SPM12, Welcome Department of Cognitive Neurology, Institute of Neurology, London, UK; https://www.fil.ion.ucl.ac.uk/spm) and carried out with the SPMMouse (http://www.spmmouse.org/) [74] animal brain toolbox implemented in MATLAB R2017b (MathWorks Inc., Natick, MA). A voxel-wise two-sample t test was used to compare the [^18^F]ROStrace SUVR_40-60_ parametric images between WT and APP/PS1 in each age group (5, 10, and 16 mo.) for both female and male. Due to the small sample size of the studies, the SPMMouse analyses were evaluated by using a relative stringent threshold of a p value < 0.005 with uncorrected statistic and a voxel extent of 50.

### Ex vivo autoradiography (ARG)

Mouse brains were collected immediately after [^18^F]ROStrace micro-PET scan completion. The mouse brains were frozen and sectioned sagittally with a thickness of 30 µm on a cryostat (Leica Biosystems, Germany), then the tissue slides were air dried for next step. The slides were exposed to phosphor plates (BAS 2040, GE Healthcare, Chicago, IL, USA) for 1 day and the ARG images were digitized by the Typhoon FLA 7000 (West Avenue, Stamford, USA). The ARG images were analyzed by using Pmod software. The ROI of periaqueductal gray was manually delineated for further calculating the ratio of images relative to it.

### Immunohistochemical (IHC) analysis

Collected brain tissues were embedded with Tissue-Tek optimal cutting temperature (Sakura, Japan) and frozen, then cut into 12 µm sagittal sections. The brain sections were fixed with 4% paraformaldehyde (PFA) at room temperature before staining, and then rinsed with phosphate buffered saline (PBS). Then, 15 mins of 0.1% Triton X-100 in PBS was applied to permeabilized tissues. After rinse with PBS, brain sections were blocked with 10% bovine serum albumin (BSA) for 1 hour at room temperature. Selected slides were incubated with primary antibodies (Online Table S2) overnight at 4°C. The following day, after wash with 1% BSA in PBS, slides were incubated with corresponding secondary antibodies (Online Table S2) for 1 h at room temperature. All slides were counterstained with Hoechst (1:2000, PBS, Thermo Fisher Scientific, H3570) and mounted with VectaShield Antifade Mounting Media (Vector Laboratories, 101098-042). Images were acquired by using Zeiss Axio Observer Microscope and Zeiss 710 Confocal Microscope (Germany). Panoramic brain tissue images were acquired by Keyence BZ-X800 Microscope (Osaka, Japan). For cortex amyloid burden and 3NT-positive cells quantification, images were uploaded onto Fiji software, amyloid plaques and 3NT-positive cells within the ROI (1.73mm x 0.65 mm) were counted manually by using the cell counter plugin, and quantification results were scaled to n/mm^2^. The average plaque sizes of each Aβ plaque in brain was measured by Zen software (Zeiss, Germany).

### Statistical analysis

All the statistical analyses were performed on GraphPad Prism software, version 7.02 (GraphPad Inc., San Diego, CA), and the results were presented as mean ± standard error (SEM) or mean ± standard deviation (SD) as noted. The statistical significance between different animal groups was determined either with one-way or two-way ANOVA following Sidak’s or Turkey *post hoc t* test. A p value < 0.05 was considered as the threshold of statistically significant.

## Supplemental Figures

**Fig SI 1.**
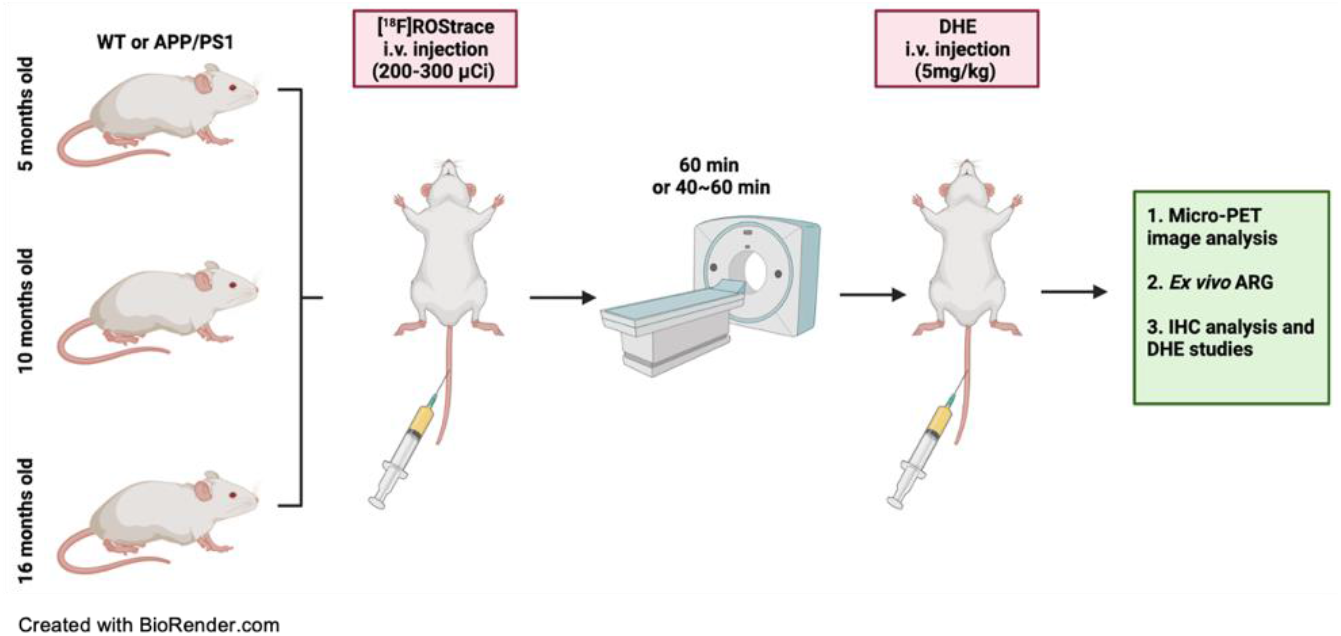
Schematic illustration of experimental scheme. DHE: dihydroethidium; ARG: autoradiography.

**Fig. SI 2.**
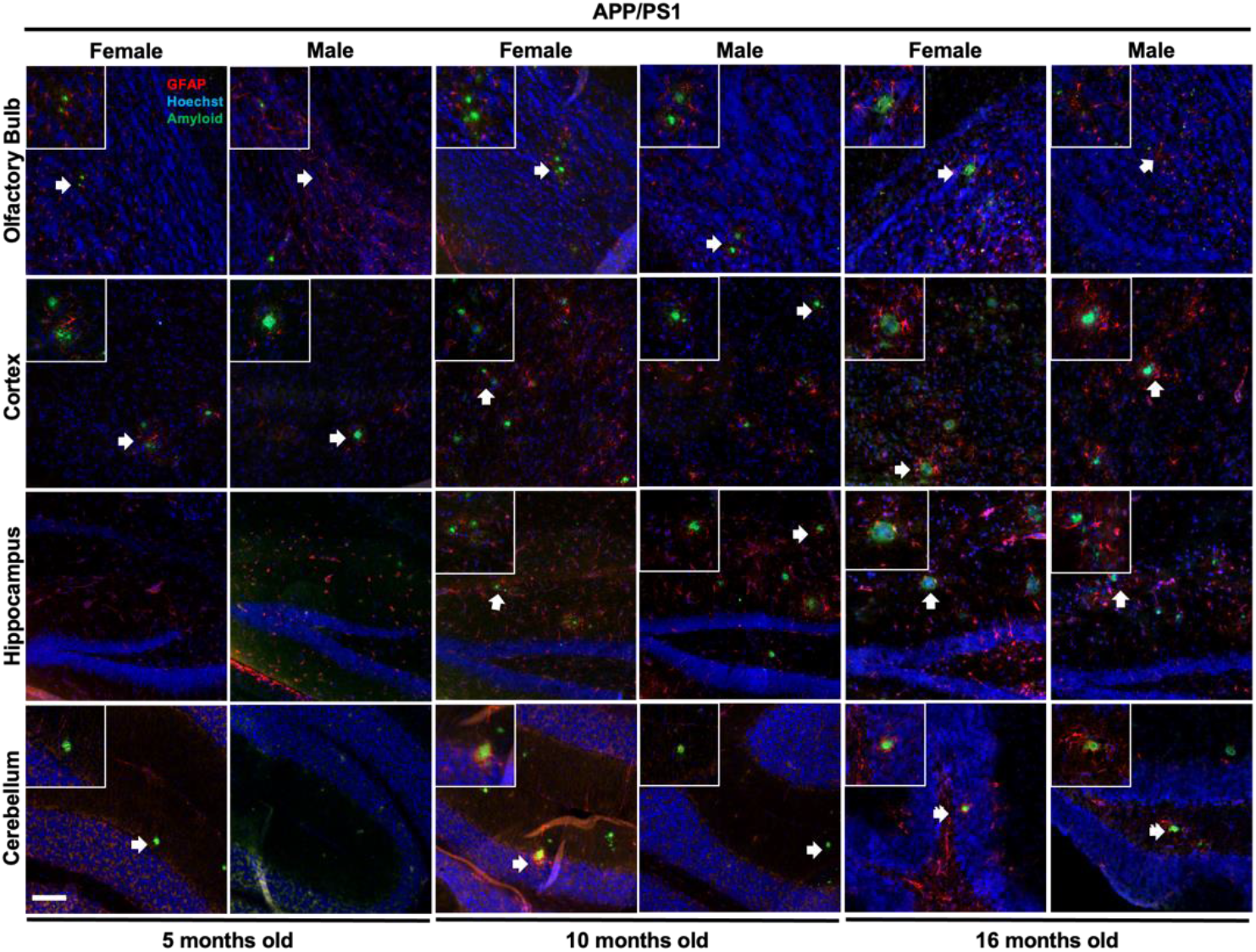
Comparison of amyloid plaques spatial distribution in age-matched APP/PS1 male and female mouse brains. Micrographs of 4 key regions from age-matched WT and APP/PS1 brain (olfactory bulbs (top row), cortex (second row), hippocampus (third row) and cerebellum (bottom row)) labeling Aβ (green), astrocytes (red), and nuclei (blue). Inlets are magnified (20x) regions of that indicated by white arrows. Scale bar = 100 μm.

**Fig. SI 3.**
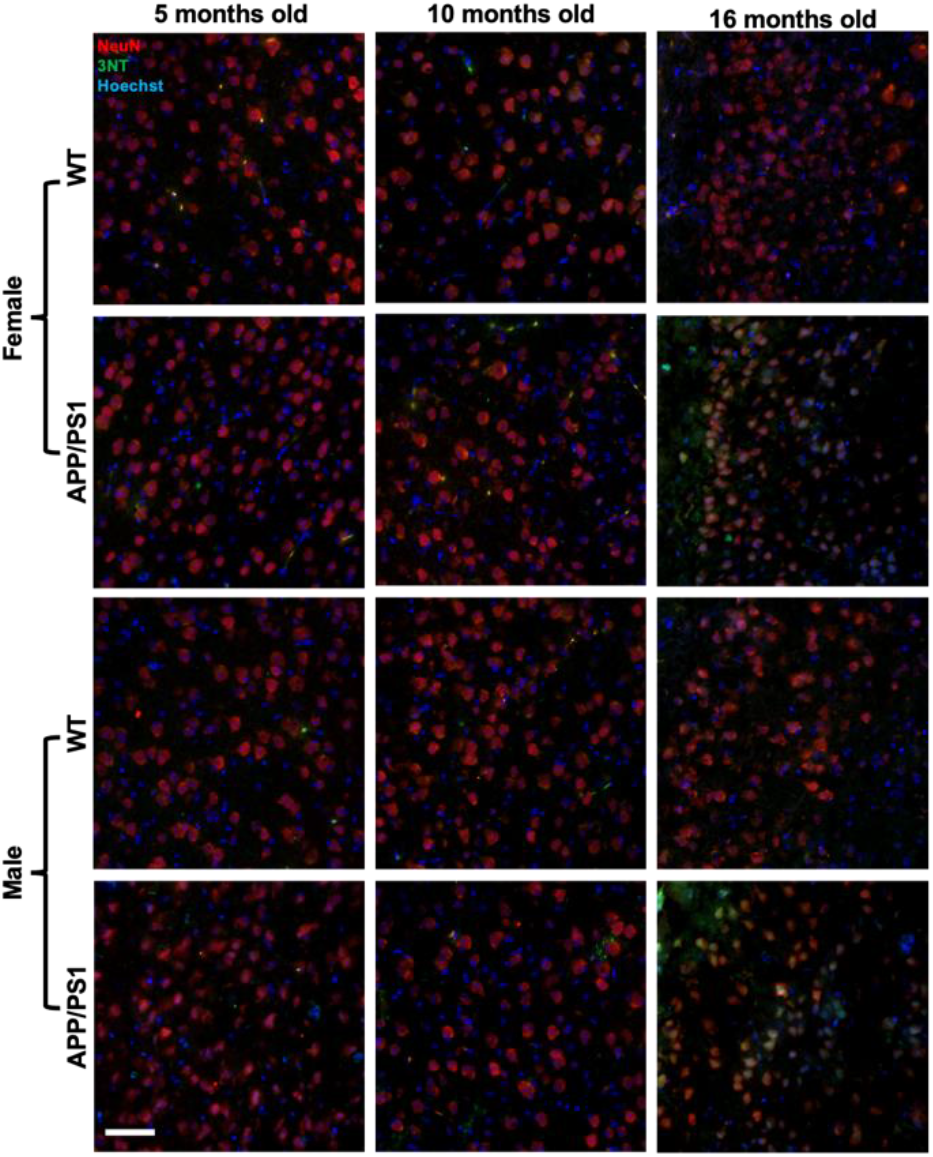
Oxidative damage detected by 3NT in neurons over time in WT and APP/PS1 cortex. Images of cortex region from age and gender-matched WT and APP/PS1 mouse brain tissues stained with anti-3NT (green) NeuN (red) and Hoechst (blue). Scale bar = 50 μm.

**Table S1.**
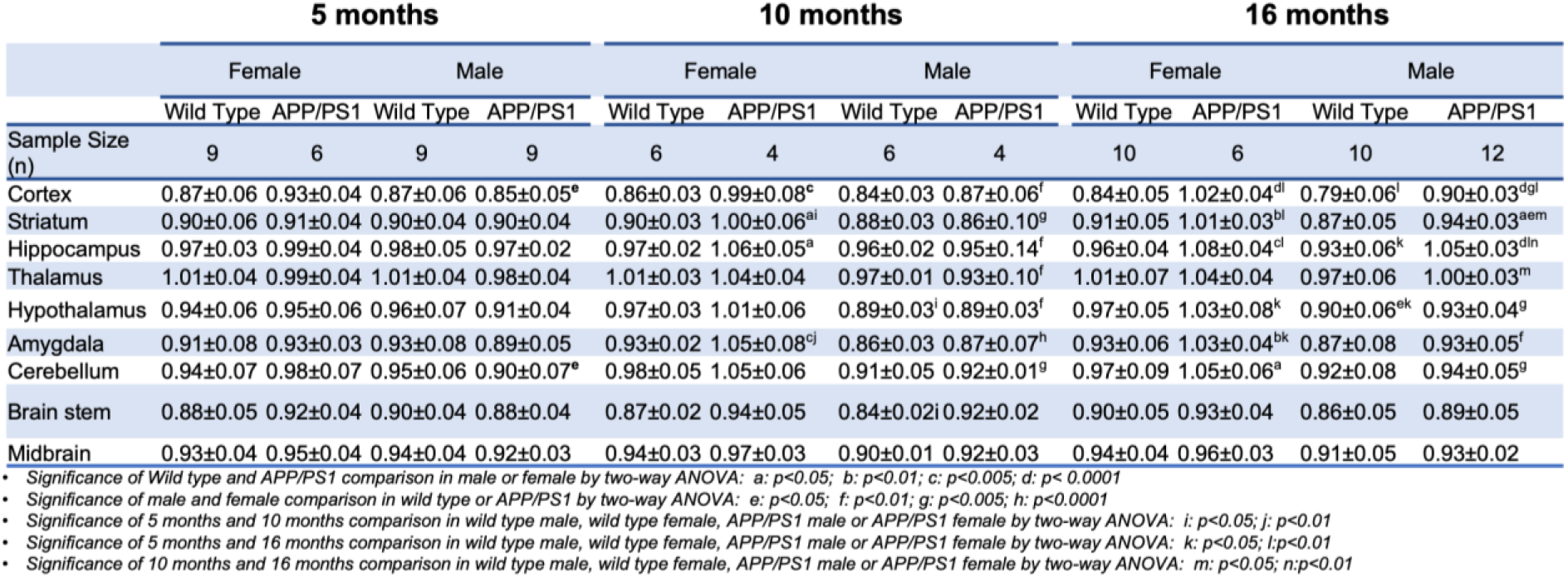
Summary of regional SUVR_40-60_ statistical analysis in age and gender-matched APP/PS1 and WT mice at the age of 5, 10, and 16 mo.

**Table S2.** Information of antibodies

| Name | Species | Cat number | Supplier | Application |
| --- | --- | --- | --- | --- |
| Anti-Glial Fibrillary Acidic Protein (GFAP) | Rabbit | AB5804 | Millipore | IF: 1:200 in 1% BSA in PBS |
| Anti-Nitrotyrosine | Rabbit | AB5411 | Millipore | IF: 1:100 in 1% BSA in PBS |
| Anti-beta Amyloid 1-42 | Rabbit | AB180956 | Abcam | IF: 1:200 in 1% BSA in PBS |
| IBA-1 | Goat | AB5076 | Abcam | IF: 1:200 in 1% BSA in PBS |
| IBA-1 | Rabbit | 019-19741 | Wako | IF: 1:200 in 1% BSA in PBS |
| NeuN | Rabbit | ab104224 | Abcam | IF: 1:200 in 1% BSA in PBS |
| Alexa Fluor 488 | Goat | A11008 | Thermo Fisher Scientific | IF: 1:200 or 1:300 in 1% BSA in PBS |
| Alexa Fluor 568 | Goat | A11011 | Thermo Fisher Scientific | IF: 1:300 in 1% BSA in PBS |
| Alexa Fluor 647 | Goat | A21235 | Thermo Fisher Scientific | IF: 1:300 in 1% BSA in PBS |
| Alexa Fluor 594 | Donkey | A11058 | Thermo Fisher Scientific | IF: 1:300 in 1% BSA in PBS |

## Acknowledgments

We thank Eric Blankemeyer and the University of Pennsylvania Small Animal Imaging Facility for the assistance with micro-PET image studies.

## Statements and Declarations

### Author Contributions

The study was designed by Robert H. Mach, Meagan J. McManus, Catherine Hou, Yi Zhu and Chia-Ju Hsieh. Material preparation, data collection, and analysis were performed by Chia-Ju Hsieh, Catherine Hou, Yi Zhu, Ji Youn Lee, Neha Kohli, Evan Gallagher, Kuiying Xu, Hsiaoju Lee, and Shihong Li. The manuscript was written by Meagan J. McManus, Chia-Ju Hsieh, and Robert H. Mach. All authors read and approved the final manuscript.

### Ethics Approval

All procedures performed in studies involving animals were in accordance with the ethical standards of the University of Pennsylvania Institutional Animal Care and Use Committee (IACUC). This article does not contain any studies with human participants performed by any of the authors.

